# Tissue specific auxin biosynthesis regulates leaf vein patterning

**DOI:** 10.1101/184275

**Authors:** Irina Kneuper, William Teale, Jonathan Edward Dawson, Ryuji Tsugeki, Klaus Palme, Eleni Katifori, Franck Anicet Ditengou

## Abstract

The plant hormone auxin (indole-3-acetic acid, IAA) has a profound influence over plant cell growth and differentiation. Current understanding of vein development in leaves is based on the canalization of auxin into self-reinforcing streams which determine the sites of vascular cell differentiation. However, the role of auxin biosynthesis during leaf development in the context of leaf vein patterning has not been much studied so far. Here we characterize the context specific importance of auxin biosynthesis, auxin transport and mechanical regulations in a growing leaf. We show that domains of auxin biosynthesis predict the positioning of vascular cells. In mutants that have reduced capacity in auxin biosynthesis, leaf vein formation is decreased. While exogenous application of auxin does not compensate the loss of vein formation in auxin biosynthesis mutants, inhibition of polar auxin transport does compensate the vein-less phenotype, suggesting that the site-specific accumulation of auxin, which is likely to be mainly caused by the local auxin biosynthesis, is important for leaf vein formation. Our computational model of midvein development brings forth the interplay of cell stiffness and auxin dependent cell division. We propose that local auxin biosynthesis has the integral role in leaf vascular development.

**Highlights:**

- Built spatially and temporally resolved auxin biosynthesis map in growing leaf primordium of Arabidopsis.
- Expression domains of auxin biosynthetic enzymes within primordia strongly correlated with leaf vein initiation.
- Results show that domains of auxin biosynthesis within primordia drive leaf vein initiation and patterning.

**Highlights and eTOC Blurb:** Using modelling and a spatiotemporal analysis of auxin biosynthesis and transport, Kneuper et al. show that tissue specific auxin biosynthesis defines places of vein initiation hence underlining the importance of auxin concentration in vein initiation.

## Introduction

Vascular systems are continuous networks of cells which connect a wide range of tissues. In leaves, veins form characteristic patterns which support photosynthesis in the surrounding mesophyll cells. However, despite leaf venation patterning being important for the overall fitness of the plant (Blonder et al., 2011), the processes which guide vein placement are not well understood. In plants the eventual pattern of the vascular network is widely believed to be controlled by the phytohormone auxin (Bruck and Paolillo, 1984), via its canalization into self-reinforcing streams (Effendi et al., 2015; Sachs, 1969). Support for with-the-flux-based explanations of leaf vascular development followed the identification of PIN1: a long-predicted auxin efflux protein and the earliest marker of leaf vascular cell identity (Galweiler et al., 1998; Scarpella et al., 2006). The polar localization of PIN1 in leaf epidermal cells has offered a mechanism which focuses auxin to specific epidermal domains which correlate with the directions of future vein development (Scarpella *et al*. 2006).

Experimental studies on auxin canalization have been corroborated with theoretical models based on the auxin canalization hypothesis, that have shown how reinforcing fluxes in a fixed domain of leaf tissue can create channels of auxin that resemble the vein patterns observed in leaves (Rolland-Lagan and Prusinkiewicz, 2005). However, the robustness of canalization models is reduced when applied to a growing tissue (Feller et al., 2015; Lee et al., 2014). The canalization hypothesis states that the position of auxin sources in the leaf primordium is critical for vein formation. It has been proposed that these sources are derived from the focal points of epidermal auxin flux, from where auxin flows in channels towards the base of the leaf (Scarpella et al., 2006). However, in addition to epidermis-derived auxin (Abley et al., 2016), auxin is also synthesized in the lamina of the growing primordium; an observation which has, to date, not been considered by any models. In support of a major role for auxin biosynthesis in vascular development in leaves, Cheng and colleagues observed progressively fewer and more discontinuous vascular strands when less auxin is synthesized in the primordium (Cheng et al., 2006), suggesting that local biosynthesis may lift auxin concentration above certain thresholds required for vascular tissues to form.

In this study, which combines a series of experiments with theory and assesses the role of auxin biosynthesis in vein initiation and patterning in the growing leaf, we find that leaf vein initiation is strongly correlated with the expression domains of auxin biosynthetic enzymes within primordia. We build an auxin-transport-independent model of vascular tissue development, and use it to identify cell expansion, auxin production rate, cellular auxin concentration and the auxin concentration-dependent cell growth of non-auxin producing cells as a robust, minimal set of parameters, for spontaneous organization of primordium cells into a midvein.

## Results

Reducing a plant’s capacity to synthesize auxin, either genetically or by the application of enzyme inhibitors, impairs the development of leaf vein networks (Nishimura et al., 2014; Stepanova et al., 2008). In mutants defective in auxin biosynthesis, leaves have fewer, more widely spaced veins which are often disorganized and discontinuous (Figures 1A and S1A-S1E, S1H). We therefore first wanted to ascertain whether auxin biosynthesis acts exclusively through increasing the overall amount of auxin available for transport-based patterning mechanisms. However, exogenous application of auxin cannot rescue aberrant vein development in *wei8-1tar2-1*, a mutant defective in auxin biosynthesis (Stepanova et al., 2011) (Figure 1), suggesting that it is not simply the amount of auxin produced which is important, but that the site of auxin production is also important for vein formation.

**Figure 1.**
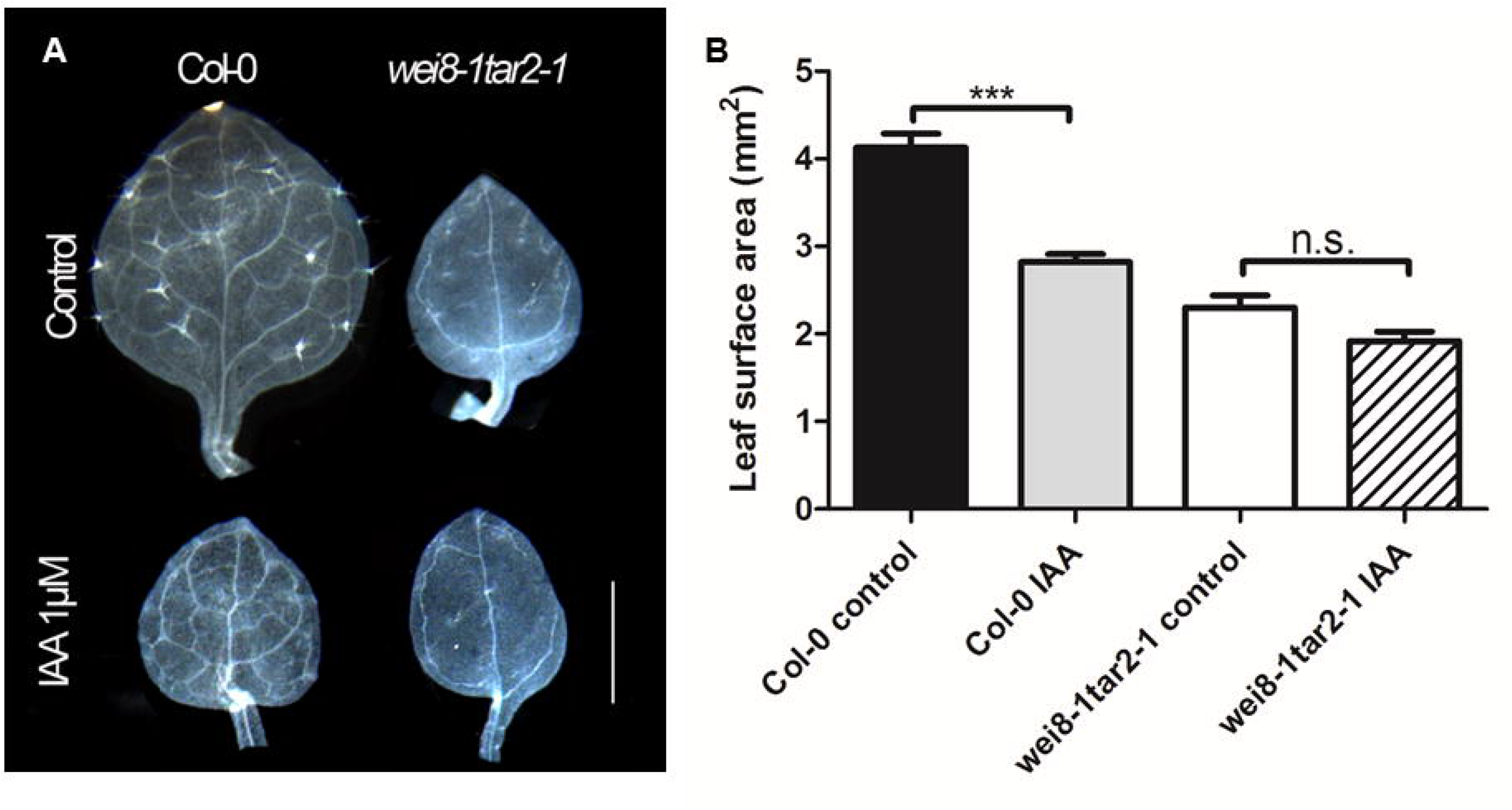
Auxin biosynthesis mutants treated with auxin. (A) Vein patterning in 12 day-old leaves WT (Col-0) and auxin biosynthesis mutant *wei8-1 tar2-1* grown in presence or absence of 1 μM IAA. (B) Quantification of leaf surface area of plants presented in (A). Asterisk (***) indicates significant difference at P<0.001 (T-test). Data are mean (*n*=40 ± s.e.). n.s., non-significant difference at P<0.05 (T-test). Scale Bar, 1mm.

Two families of enzymes act in series to catalyze the indole-3-pyruvic acid (IPA)-dependent production of IAA: tryptophan aminotransferases (TAA1/WEI8, TAR1 and TAR2) and YUCCA flavin monooxygenases (Mashiguchi et al., 2011; Stepanova et al., 2011; Zhao, 2012). Here, we focused on YUC1, YUC2, YUC4, YUC6, as they are the main YUCs for auxin biosynthesis in shoots, whereas the other YUCs (i.e. YUC3, YUC5, YUC7, YUC8, YUC9) are responsible for producing auxin in roots (Won et al., 2011; Zhao, 2012). In order to resolve spatial synthesis of auxin within the lamina of a growing leaf, we visualized the expression pattern of auxin biosynthetic genes and their correlation with pre-provascular cells in the leaf primordium. We built an auxin-biosynthesis map in Arabidopsis, at cellular resolution using well established standard techniques (imaging of GFP fusion proteins, promoter GUS fusions), which reflect faithfully the expression of auxin biosynthetic genes in leaf primordia over time. At between 2 and 5 days after germination (DAG), enzymes of YUCCA and TAA1 families showed highly localized expression patterns in which the future sites of vascular cells can be clearly seen (Figures 2; S2 and S3). Over this time period, expression domains of both classes of enzymes were restricted to the regions of leaf primordia in which veins are formed, and the border between the abaxial and adaxial sides (lower panel in Figure 2A and Figure S3). TAA-type aminotransferases and *YUCCA* genes are expressed in distinct as well as in overlapping domains (Figures 2, S2 and S3). Note particularly the overlapping expression of TAA1, TAR2, and YUC4 during the early stage of midvein development in 2 day-old primordia (Figures 2, S2 and S3). Since TAA and YUC act in succession in the same IPA pathway to produce IAA from tryptophan, co-expression of TAA and YUC in the same cells increases the chance to produce IAA.

**Figure 2.**
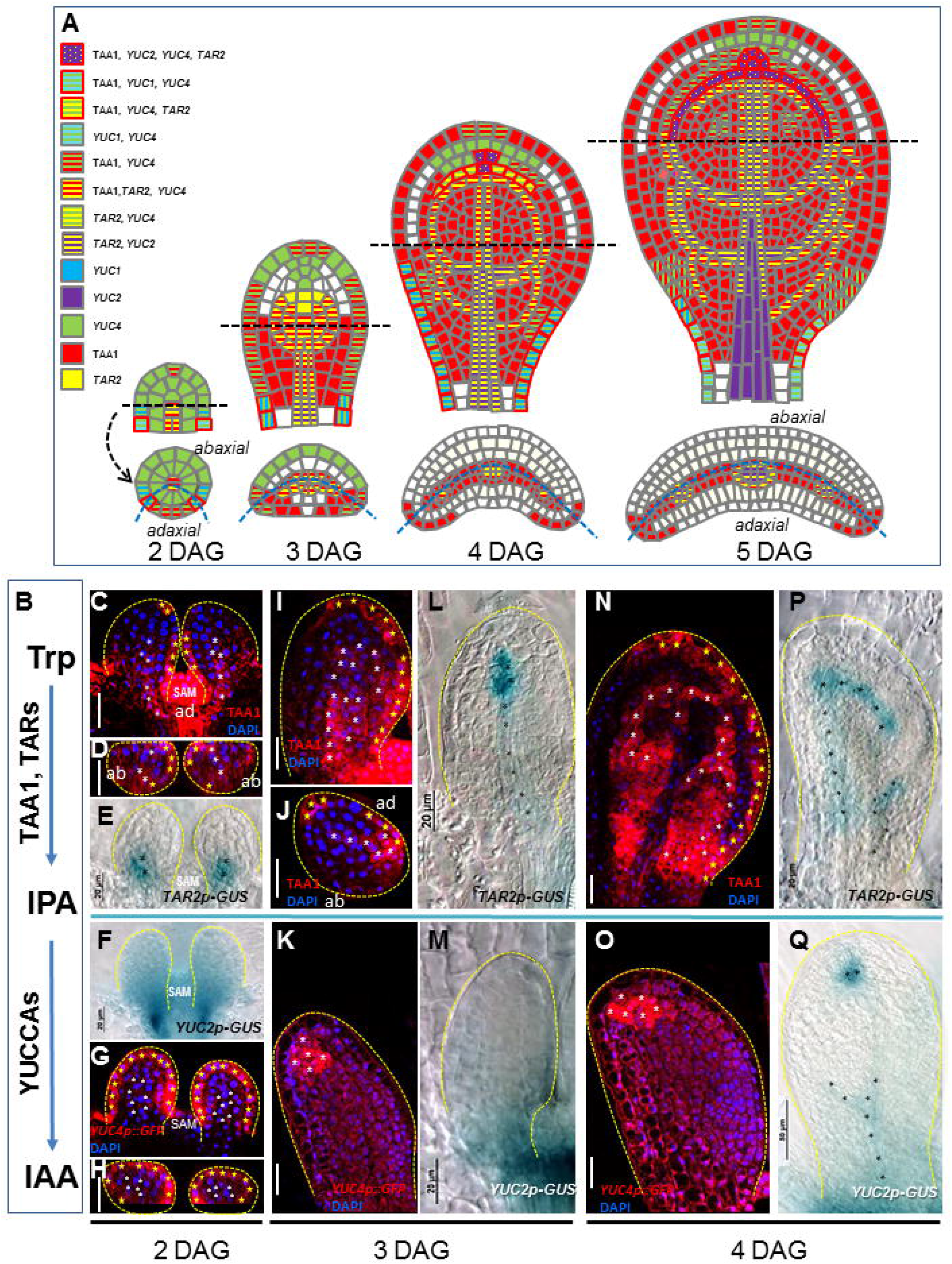
Time resolved auxin biosynthesis map in Arabidopsis leaf. Combinatorial expression of auxin biosynthetic genes (*TAA1, TAR2, YUC1, YUC2, YUC4*) at 2 DAG (C-H), 3 DAG (I-M) and 4 DAG (N-Q). (A) Longitudinal (above) and transverse (below) leaf sections. The blue dashed line defines the separation between adaxial (ad) and abaxial (ab) leaf polarity. (B) Schematic representation of the major auxin biosynthesis pathway (Trp, tryptophan; IPA, indole 3-pyruvic acid; IAA, indole3-acetic acid). (C-Q) Time resolved expression pattern of *pTAA1::TAA1-GFP* (C, D, I, J, N), *TAR2::GUS* (E, L, P), YUC2::GUS (F, M, Q), *YUC4::GFP* (G, H, K, O). D, H, and J are optical transversal sections of the leaf. Asterisks: yellow asterisks indicate TAA1 and *YUC4* expression in the epidermis; white asterisks indicate TAA1 and YUC4 expression in the lamina; black asterisks show TAR2 and YUC2 expression in provascular tissues. SAM indicates the position of the shoot apical meristem. Scale bar, 20 μm unless otherwise indicated. DAG, days after germination. The data supporting auxin biosynthesis in 5 DAG leaf are presented in Figures S1 and S2.

The effect of inhibiting polar auxin transport on vein patterning, either by introducing genetic lesions (Okada et al., 1991), or by the exogenous application of diverse auxin transport inhibitors (Mattsson et al., 1999; Sieburth, 1999), has been known for decades. The inhibition of auxin transport, either genetically or by the application of inhibitors, leads to the formation of small, round leaves with an indistinct midvein and fused vascular bundles around the periphery of their distal end (Mattsson et al., 2003; Mattsson et al., 1999). Therefore, it was proposed that in a developing Arabidopsis leaf, auxin transport defines the localization of leaf veins (Scarpella et al., 2006). In order to ascertain whether auxin transport also defines sites of auxin biosynthesis, we examined the distribution of cells which contained auxin biosynthetic enzymes (or their transcripts) in the leaf primordium after inhibition of auxin efflux.

In Arabidopsis, 1-N-naphthylphthalamic acid (NPA) abolishes polar auxin transport at concentrations above 10 μM (Thomson et al., 1973). After treatment with increasing concentrations of NPA up to 320 μM, leaves became round and gradually smaller (Figure 3A). Furthermore, they developed an increased number of secondary veins and an exaggerated mesh of fused tracheary elements which surrounded the leaf margin, when compared to leaves of untreated plants (Cai et al., 2014) (Figure 3A). After blocking auxin transport, the domains of auxin biosynthetic enzymes expanded and continued to predict the regions of vein initiation (Figures 3B-3E, S4). In leaf lamina, with or without NPA treatment, PIN1 expression in the lamina was confined to cells which lay within auxin biosynthetic domains (Figure 3C-3F). The change in shape of the leaf after NPA treatment is correlated with the proliferation of provascular cells expressing both PIN1 and auxin biosynthetic genes in these leaves (Figure 3B-3F). In summary, NPA application changed the auxin biosynthesis pattern, but veins were nevertheless still formed in regions of auxin biosynthesis [Fig. S4A (lower panel), S4B and S4C]. Therefore, blocking auxin efflux causes auxin levels in non-vascular cells to drop, which then lowers cell division rates in the non-vascular cells causing vascular cells to proliferate laterally (Figure 3B-3F). NPA also induced the accumulation of auxin at the distal end of the leaf (as indicated by DR5::Venus in Figure 3H, compare upper versus lower panel), suggesting the distal sites also produce auxin (Abley et al., 2016; Avsian-Kretchmer et al., 2002).

**Figure 3.**
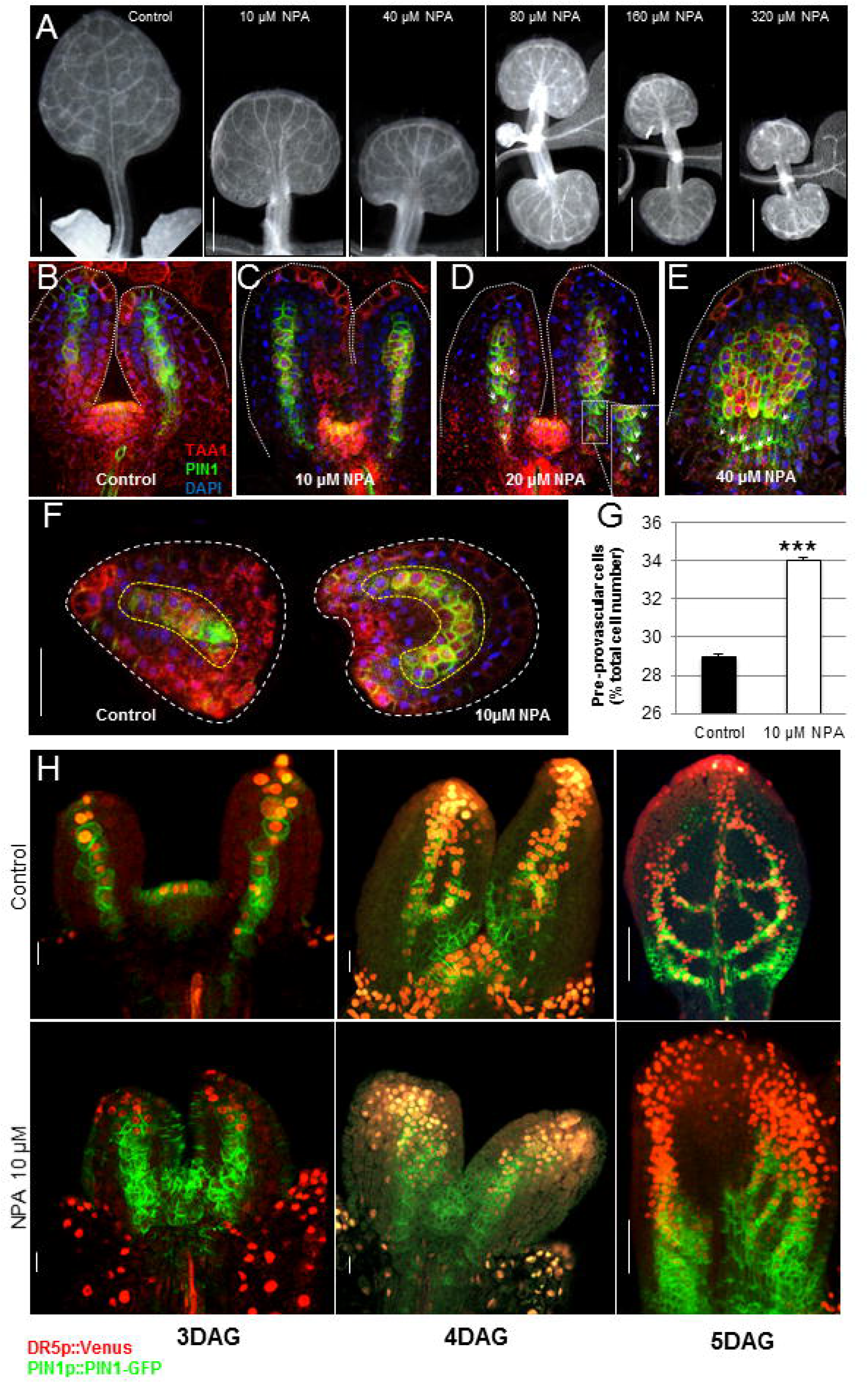
Effect of NPA on leaf vein patterning, auxin biosynthesis and distribution. (A) Vein patterning in 12 day-old leaves grown in presence of increasing concentrations of NPA. (B-F), Immunodetection of both TAA1-GFP and PIN1 in 4 day-old leaves grown in presence of NPA. Boxed Inset in (D) shows polar PIN1 at the plasma membrane of vascular cells arrowed in (D) and (E). F, Transversal sections of leaves presented in B and C, respectively. White dashed lines indicate leaf boundaries. Yellow dashed lines indicate pre-provascular cells expressing both TAA1 (red) and PIN1 (green). Note the proliferation of pre-provascular cells in a concave shape in 10 μM treated leaf. G, Percentage of cells expressing both PIN1 and TAA1 in leaf lamina. By dividing by the total number of cells constituting leaf primordium. Stars (*) indicate significant difference to control at P<0.01 (t-test). Data are means (n=40 ± s.e.). H, DR5::Venus (auxin levels) and PIN1p::PIN1-GFP in 3, 4 and 5 day-old leaves. Top panel: control leaves. Lower panel: leaves treated with 10 μM NPA. Scale, 500 μm in (A), 200 μm (F) and 10 μm (H).

### *In silico* modeling of spontaneous patterning of vascular cells in the leaf primordium

In order to test whether cell specific auxin biosynthesis and its impact on neighboring cells is able to drive leaf vein patterning, we developed a theoretical model and performed a series of *in silico* experiments. We adapted Virtual Leaf, an open-source cell-based modeling framework that describes cells of the leaf lamina as a two-dimensional layer of interconnected polygons and accounts for mechanical properties of the tissue (Merks et al., 2011). Cell growth (an irreversible increase in cell area) proceeds in a quasi-static way at a rate which is defined by cell turgor pressure, a feature that can be influenced by auxin (see Methods section). A cell divides each time its area doubles, which invariably results in tissue growth over time. Cell wall stiffness and mechanics are included in the model by the definition of a resting length for each cell wall element. The shape of the tissue at each stage of growth is determined by minimizing the generalized energy function that contains a cell area term and a cell wall elasticity term (see computational model description in the Methods section).

To model leaf vein patterning in a growing leaf, we created a leaf tissue template that closely resembles a 2-day-old leaf primordium as our initial condition (Figure 4A). Following Lee, Feugier *et al*. (2014) and Merks, Guravage *et al*. (2011), we modelled primary vein development in two dimensions. The model tissue template consisted of cells expressing auxin biosynthesis genes, labeled as auxin producer cells, and cells that do not synthesize auxin. In the model, auxin producer cells correspond to the cells that express the auxin biosynthetic gene *TAR2*, and their layout in the tissue resembles the *TAR2*-driven GUS expression in 2-day-old control leaf-primordium (Figure 4A). The mechanical constraint due to the attachment of the leaf to the meristem is modeled by including a row of cells at the proximal part of the leaf (the petiole). These cells also act as auxin sinks, to model auxin drainage from the leaf via established veins at the stem (see computational model description in the Methods section). Further assumptions of our model are as follows: i) inter-celullar auxin transport via a diffusion-like process (PIN1 has a non-polar distribution in pre-provascular cells), ii) every cell in the leaf lamina grows by increasing its area at the same rate, iii) auxin is known to promote cell expansion in aerial tissues (Fendrych et al., 2016; Perrot-Rechenmann, 2010), therefore when auxin concentration in a non-auxin producing cell crosses a threshold, then this cell starts expanding at a higher rate, iv) auxin-synthesising cells grow at all times at the basic rate (as defined in (ii)), and v) a relatively stiff epidermal cell layer, as it is known that leaf epidermal cells are stiffer than laminar cells (Onoda et al., 2015).

**Figure 4.**
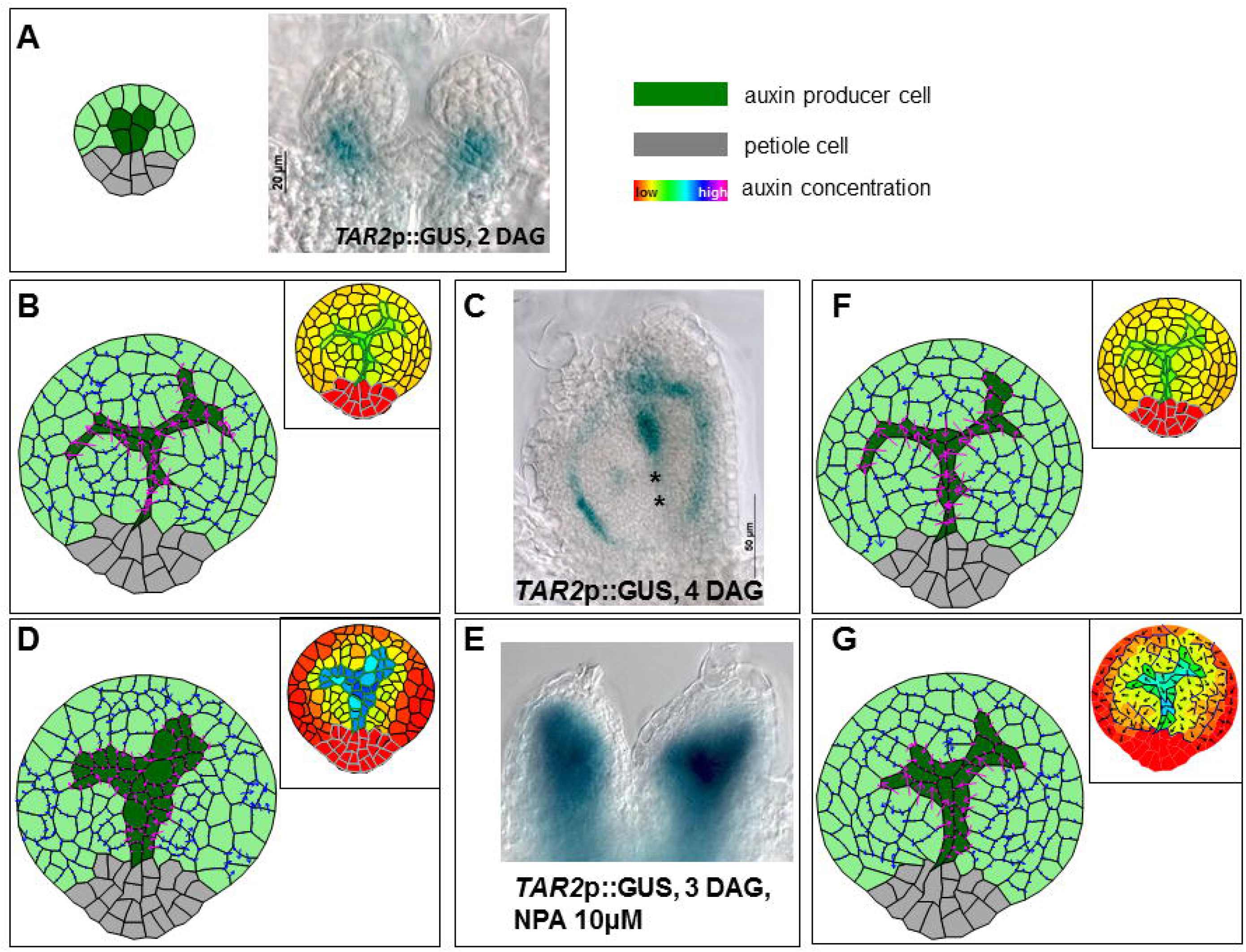
*In silico* modeling of spontaneous patterning of vascular tissue in leaf primordium. (A) Left, leaf template resembling a leaf primordium. The model assumes local auxin synthesis (dark green colored cells), leaf petiole cells (grey colored cells), a relatively stiff outermost surface of the epidermal layer, non-directional transport of auxin and auxin-dependent cell growth; right *TAR2*-driven GUS expression in 2-day-old control leaf-primordia. (B) Left, simulation result of our model with the parameter values as given in Table S1. The model was able to reproduce a realistic mid-vein made up of elongated cells and branched vasculature, as observed in 4-day-old control leaf-primordia (see C). Forces on the vertices of a cell in the leaf tissue is shown by an arrow representing the force vector, in the case of a control-leaf; Magenta colored arrows represent forces on the vertices of midvein cells, and blue arrows represent forces on the vertices of other cells in the leaf primordia. Inset shows cellular auxin concentration in the leaf primordia. (C) *TAR2*-driven GUS expression in a 4-day-old control leaf-primordia. (D) Simulation result with a 25-fold lowered rate of auxin transport. All the other parameter values were kept the same as in (C). Inset shows cellular auxin concentration in the leaf primordia. (E) *TAR2*-driven GUS expression in 3-day-old NPA-treated leaf-primordia. (F) Simulation result for reduced auxin production and reduced transport rate. A realistic midvein could be produced in simulation only when the overall cell area growth rate was decreased, consistent with NPA treatment rescue of an auxin biosynthesis mutant. (G) *In silico* modeling of PIN1 based auxin transport. The model incorporates PIN1 based transport of auxin, and is able to reproduce a realistic mid-vein even in the case of no diffusion (*d=0*) and PIN1 based transport only. Inset shows cellular auxin concentration in the leaf primordia. The arrows indicate the net direction, but not the magnitude of the auxin flux. The lack of clear polarity of the auxin flow in the epidermal cells is likely due to the absence of any external source of epidermal auxin. Asterisks (*) denote provascular cells.

Simulations of this simple model, using the experimentally reported values of various parameters (see computational model description, Table S1) (Mitchison 1980), generated an *in silico* leaf-primordium that had grown to a size of either approximately 150 cells (Figures 4B) or 900 cells (Figure S5). This simulation result showed that, for a wide range of parameter values determining auxin-dependent cell growth (see computational model description in the Methods section), and without including polarized auxin transport, this model is able to reproduce i) proliferation of auxin synthesizing cells, ii) correct midvein positioning and iii) coordinated vascular cell elongation, as experimentally observed (Figures 4B-4C; S5 and Video S1). The shape of midvein generated by the model is robust with respect to different initial geometry and number of auxin synthesizing cells (Fig. 4A left and right panel and Fig. S7). Our simulations showed that localized auxin synthesis, followed by auxin diffusion, was able to account for the establishment of a time-dependent auxin gradient across the tissue of leaf primordium (inset of Figure 4B). Such an auxin gradient caused cells to grow at different growth rates. This in turn induced transverse forces acting laterally on the walls of midvein cells. These forces are are much larger in magnitude than the forces acting on other cells in the leaf lamina (Figure 4B). These forces exerted by the neighboring cells on the midvein cells, resulted in non-trivial strain and force distributions in the tissue that prevented the midvein cells from proliferating and thus resulted in the development of a thin vascular strand of elongated cells as is characteristic of procambial cells (Scarpella et al., 2006).

We next studied *in silico*, the impact of reducing auxin transport rates on midvein formation. An NPA-dependent reduction in auxin efflux led to a change in leaf vascular patterning and an expansion of-the *TAR2* expressing domain, suggesting an extension of auxin biosynthesis sites (Figure 4E). Simulations of our model showed that a 25-fold lowering of the rate of auxin transport altered the formation of the midvein, and resulted in much wider proliferation of auxin-producing cells, as observed in experiments (Figure 4D). Reducing auxin transport rate led to higher auxin concentrations in auxin producer cells and steeper auxin gradients in the primordia (Figure 3H lower panel and inset of Figure 4D), with non-auxin producing cell receiving less auxin from auxin producing cells and hence, expanding in that case less than in leaves not treated with NPA. As a result, the auxin-dependent forces exerted by the neighboring cells on the midvein cells were not strong enough to prevent the proliferation of auxin producer cells (Figure 4D). The model predicted a midvein, several cells wide with a high auxin concentration, just as was observed in plants grown on inhibitory concentrations of NPA (Figures 3C-3E, 3F-3G, 3H and 4E). Furthermore, the midvein cells of the “NPA-treated” model leaf did not elongate in the same way as “untreated” simulations (Figure 4D and inset, and Video S2).

### Compensation of vein patterning in auxin biosynthesis mutants by NPA

The model predicted that NPA effects were due to the accumulation of auxin in developing vascular cells. We therefore predicted that the reduced-efflux phenotype would be rescued by simultaneously decreasing the rate of auxin synthesis. Accordingly, we simulated the treatment of auxin biosynthetic mutants with NPA by simultaneously reducing both the auxin transport rate and the auxin production rate (2-fold). Under a range of conditions, wild-type leaf vascular patterning was restored, but only when the overall cell area growth rate was decreased (Figures 4F, S6A-D) (Methods section). *In planta*, the sequential removal of auxin biosynthetic enzymes resulted in the differentiation of fewer vascular strands, or even in the complete inhibition of vein development (Figures 5C, left panel and S1A-E). However, as predicted by the model, many of these phenotypes were abrogated by the application of NPA (Figure 5C and 5D). This observation suggests that the cellular auxin concentration *per se* can define vascular cell differentiation in leaf primordia. We therefore conclude that an underlying pattern of auxin biosynthesis, influenced by the physical environment of the primordium, can direct vascular cell initiation.

**Figure 5.**
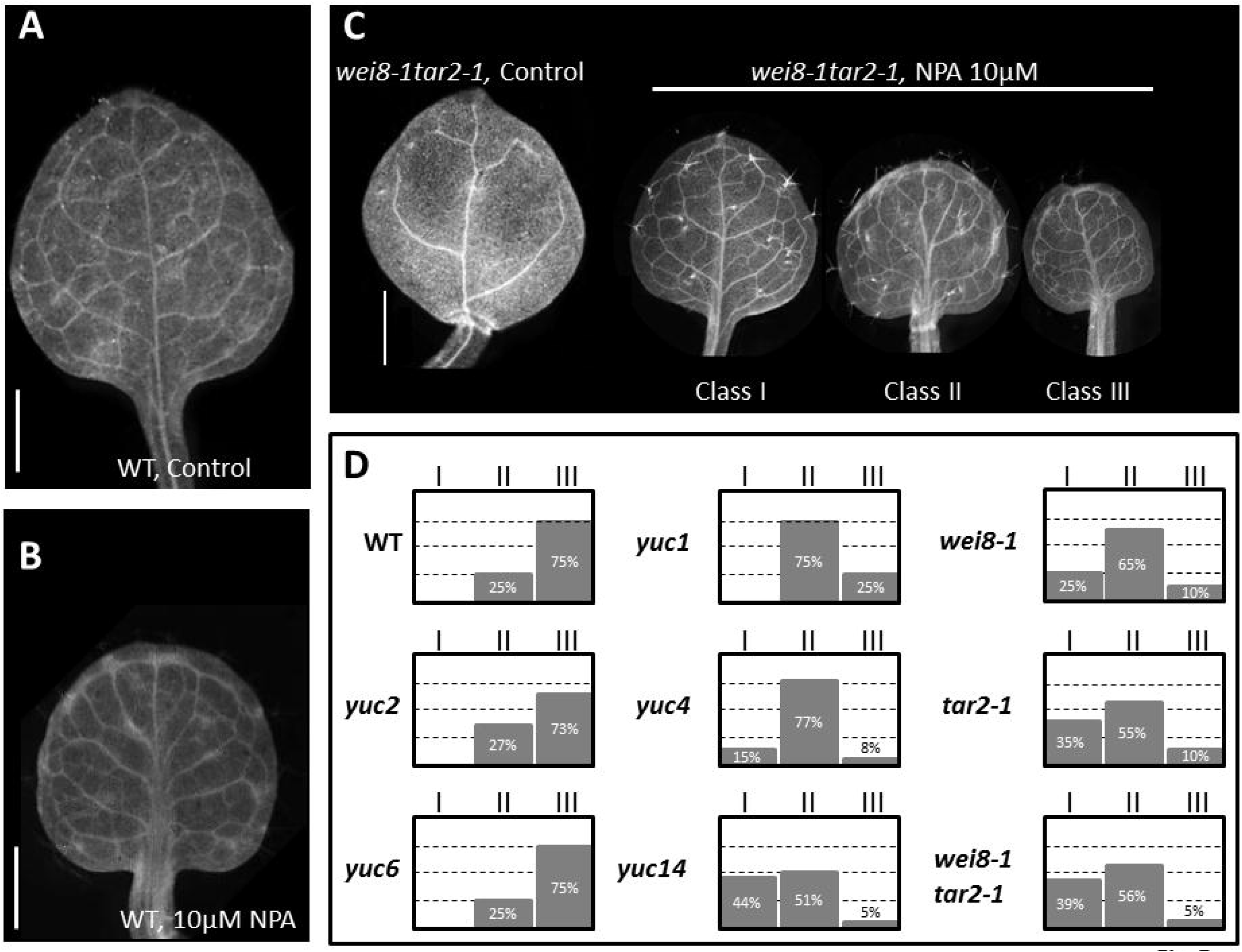
Compensation of vein patterning in auxin biosynthesis mutants by NPA. (A) Vein patterning in 10-day old wild-type (WT) leaf. (B) WT leaf after NPA treatment. (C) *wei8-1tar2-1* leaf treated with (right) or without (left) 10 μM NPA. Leaves were classified according to their vein architecture. Class I, indistinguishable from untreated WT. Leaves contain a single midvein reaching distally to the tip of the leaf. Class II, fused midveins and Class III, fused midveins and increased frequency of fused high order veins. (D) Genotype dependent distribution of Class I, II and III leaves in seedlings treated with NPA. Scale bar, 500 μm.

### *In silico* modeling of PIN1 based auxin transport

Polar auxin transport has long been considered a major driver of leaf vascular cell differentiation (Figure S1F) (Scheres and Xu, 2006). “With the flux” models of leaf vascular patterning assume that cells sense the overall rate of auxin flux and respond by reinforcing this flux with the polar deposition of PIN proteins. We therefore studied computationally the effect of polar auxin transport on the formation of the midvein in the presence of localized auxin biosynthesis (see Methods section) (Figure 4G). Our simulation results show that polar auxin transport reproduces the midvein in much the same way as in the diffusion-only case (discussed above; Figure 4B), suggesting that these two patterning mechanisms may be linked or working in parallel.

### Regulation of auxin biosynthesis by auxin

As the canalization of auxin requires a positive feedback loop between the plasma membrane’s permeability to auxin and the rate of auxin flux (Mitchison, 1981), we next addressed whether auxin biosynthesis and flux participate at a transcriptional level in such a positive feedback loop. Mutants defective in auxin biosynthesis were examined for PIN1 expression. However, the polarization of PIN1 proceeded as normal in five-day old leaves of *wei8-1tar2-1* and *yuc1yuc4* loss-of-function plants, despite a drastic reduction in the density of high order veins being observed (Figures 6B and 6C, lower panel of S1B, S1E and S1H). Next, neither TAA/TARs nor *YUCCA* transcription is induced by IAA (Figure 6D). This experiment was performed three times with similar results. Verification in Genvestigator (www.genevestigator.com), also confirmed that the expression of *TAA1, TAR1, TAR2, YUC1, YUC2, YUC4* and *YUC6* genes is not induced by auxin. Further, our results also indicate that *YUC1, YUC2, YUC4* and *YUC6* are repressed by auxin; similar results were reported by Suzuki and colleagues (Suzuki et al., 2015). Finally, we found that TAA1 expression was reduced as PIN1 accumulated and polarized in provascular cells (Figure 7B, 7C). IAA biosynthesis is therefore not induced by the canalization of IAA, but is likely to be integrated into an independent upstream patterning mechanism, as although TAA1 was seen in domains in which PIN1 is absent (Figure 7D-7I); the opposite case was never observed. These observations suggest that auxin biosynthesis is not part of the positive feedback loop proposed to initiate vasculature development in leaves.

**Figure 6.**
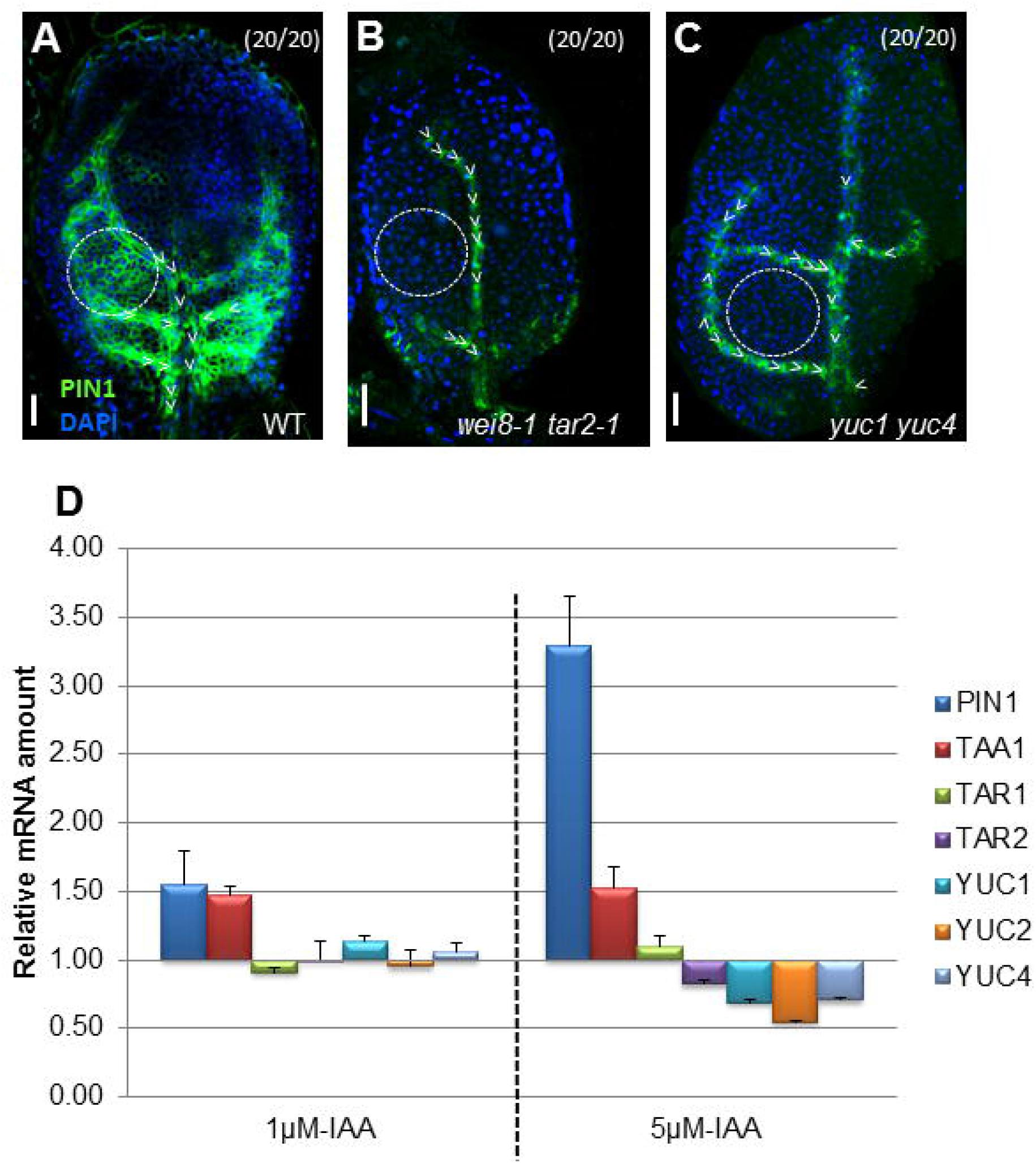
Auxin does not regulate PIN1 polarisation nor its own biosynthesis. (A), (B) and (C) Immunocytodetection of PIN1 in WT (A), in *wei8-1 tar2-1* (B) and in *yuc1 yuc4* double mutants (C). Arrowheads indicate PIN1 polarity; dashed circle, intervascular pre-provascular cells expressing PIN1 visible in WT but absent in *wei8-1 tar2-1* and in *yuc1 yuc4* double mutants. (D), Quantification of *PIN1, TAA1, TAR1, TAR2, YUC1, YUC2, YUC4* mRNA in 3 day-old WT seedlings incubated in the presence of 1 μM or 5 μM IAA for 4 hours. (Parentheses) Fraction of primordia showing the displayed features. Scale bar, 20 μm.

**Figure 7.**
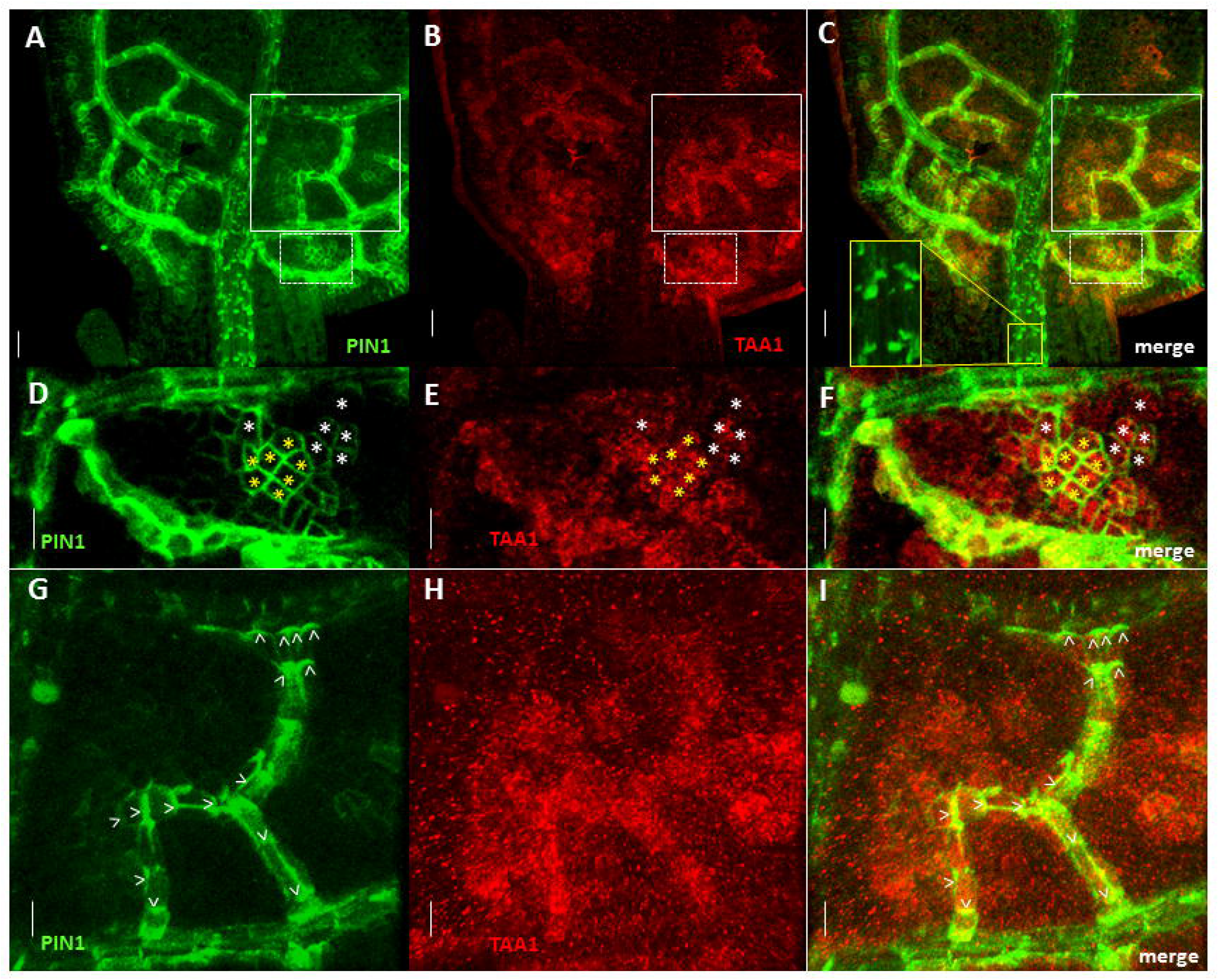
Gradual subcellular localization of PIN1 in dynamic TAA1 expression domains. PIN1 and TAA1-GFP subcellular localization in 6-day old leaf. (A-C), Dynamic expression pattern of TAA1-GFP and subcellular localization of PIN1 in pre-provascular, provascular and mature vascular cells. (A), PIN1 localization. (B), TAA1-GFP. (C), Panel (A) and panel (B) merged. (D-F), Higher magnification of images in dotted frames in (A-C), respectively. (G-I), Higher magnification of images in solid frames in (A-C), respectively). Arrowheads indicate PIN1 polarity in provascular cells. Asterisks (*): white asterisks indicate developing provascular tissues, yellow asterisks indicate pre-provascular cells with PIN1 non-polar. Yellow inset in (C) shows respectively polar PIN1 and TAA1 down regulation in mature vasculature. Scale bars, 20 μm (A-C), 10 μm (D-I).

## Discussion

In plants, polarized auxin transport is crucial for the initiation and regulation of several developmental programs. PIN-mediated auxin flux is widely accepted to be responsible for vascular patterning in leaves, with vascular differentiation occurring along axialized (canalized) auxin streams (Scarpella et al., 2006; Sieburth, 1999). Here we provide evidence that vein initiation also requires tissue specific local auxin biosynthesis. This supported by the fact that in mutants defective in local auxin biosynthesis, vein density is severely reduced (Cheng et al., 2006; Stepanova et al., 2008).

In leaf primordia, auxin transport does not only occur in developing veins, but also in epidermal cells (Abley et al., 2016). Blocking auxin transport results in smaller, round leaves which display a characteristic venation pattern. It was shown that cells accumulate more IAA when treated with NPA (Petersson et al., 2009; Petrasek et al., 2006). Therefore, we hypothesized that, since auxin is synthesized in both vein cells and epidermis cells, applying NPA would increase auxin concentration in both areas. This did not lead to an increase in leaf size, suggesting that auxin does not increase the rate of cell division in the epidermis. Similarly in roots, auxin induces cell division in pericycle cells, but not in epidermal cells (Himanen et al., 2002; Mähönen et al., 2014; Pacheco-Villalobos et al., 2016). Thus, in presence of NPA, epidermal cells (less sensitive to auxin) would divide less and as they surround proliferating vascular cells in the lamina, this would supply a possible reason for the change in mechanical properties of the leaf identified by our model.

It was shown that in wild-type leaves, midvein provascular cells have potential characters of midvein progenitor cells (Tsugeki et al., 2009). This observation is consistent with a hypothesis that clonal populations of vein cells originate from those cells of the primordium which already have a vascular identity. This hypothesis is supported by an experiment using the expression of CRE recombinase under the control of the heat shock promoter (Ichihashi et al., 2011) which showed that in developing leaf, clonal sectors give rise to connected veins.

The patterning of organ growth and development is constrained and directed by the cells’ physical environment, with mechanical stresses in particular influencing a broad range of developmental pathways (Sampathkumar et al., 2014). Microtubules responsively re-orientate their direction according to the stress landscape of a tissue in order to reinforce cells against the strains caused by directional growth (Hamant et al., 2008). Considering the pervasive influence of mechanical stress over plant development and patterning, it is likely that they also play a role in developing leaf primordium.

Our model is able to explain aspects of leaf development which include the production of auxin by pre-provascular cells of young leaves, the presence of vascular tissue in leaves treated with high concentrations of NPA (Mattsson et al., 1999), the relative insensitivity of vascular cell initiation to NPA when compared to a reduction in auxin biosynthesis, and the strong auxin response maxima observed in differentiating vascular cells (Bayer et al., 2009; Heisler et al., 2005). Using the model we present here, we were able to recreate faithfully four key features of vascular cell biogenesis in the absence of polar auxin transport: vascular strand formation, midvein positioning and coordinated vascular cell elongation. Our analysis suggests a critical role for the site of auxin biosynthesis in directing vascular cell differentiation. Thus, local auxin biosynthesis and a correspondingly high cellular auxin concentration need to be considered alongside directional auxin flux as an important factor defining leaf vascular cell patterning. This conclusion is supported by the requirement for context-specific auxin biosynthesis to complement multiple *yucca* mutants (as opposed to exogenous application) (Cheng et al., 2006) and the fact that relatively small reductions in auxin amounts at a whole-plant level can cause surprisingly severe auxin-deficient phenotypes (Stepanova et al., 2011).

The complete sequence of genes necessary auxin biosynthesis is expressed in the leaf lamina (Mashiguchi et al., 2011; Stepanova et al., 2011; Zhao, 2012). Besides TAA1 whose expression in pre-provascular cells extends to regions that do not make veins and then is downregulated once pre-provascular cells become provascular cells, TAR2 is expressed exclusively in regions that make veins, where YUC2 and YUC4 are also expressed. This highlights an established correlation between auxin biosynthesis and vein development: in mutants defective in auxin biosynthesis, PIN1 expression and high-order vein density decrease. In contrast, suppression of auxin transport in *pin* mutants or using chemical auxin transport inhibitors has only a marginal impact on vein initiation. This suggests that auxin biosynthesis may act upstream auxin transport and is necessary for leaf vein initiation. According to the auxin transport-based canalization hypothesis, it is the auxin flux (rate of auxin molecules going through the membrane in a polar PIN1-dependent manner) that initiates veins. On the other hand, leaf veins are still formed when polar auxin transport is chemically abolished (as when a seedling is treated with NPA, 2,3,5-triiodobenzoic acid (TIBA) or any other auxin transport inhibitor) (Carland et al., 2016), which is hard to be explained only by the auxin transport-based canalization hypothesis. Together these observations underline the crucial role of tissue specific auxin biosynthesis which may act prior auxin canalization.

Our model predicted that reducing auxin biosynthesis and cell growth rates in the presence of NPA would restore normal venation patterns in *wei*/*tar* mutants. Indeed, treating auxin biosynthesis mutants, which display both reduced auxin levels and growth rates (they develop smaller leaves) with NPA, did consistently restore wild-type like vein patterning. Since cells accumulate more IAA when treated with NPA (Keller et al., 2004; Petersson et al., 2009; Petrasek et al., 2006), the abnormal venation pattern in *wei*/*tar* mutants is probably due to cell specific decreases in auxin concentration. Therefore, the inhibition of auxin efflux by NPA would result in an increase of auxin concentration in pre-/provascular cells, which would then trigger the initiation of venation. However, since NPA also affects the shape of the leaf, it confirms that auxin transport in the epidermal cells plays an important role in maintaining leaf shape as previously proposed (Izhaki and Bowman, 2007; Scarpella et al., 2010). Taking into account cell auxin biosynthesis in the leaf lamina and auxin transport in epidermal cells in future models will enable us to understand how leaf shape and vein patterning are coordinated. In leaf lamina, auxin is transported through elongating vascular cells. However, it is not likely that auxin transport in elongating vascular cells is a prerequisite for midvein formation or vascular branching, since veins still form in presence of very high concentrations of NPA. Instead, the model predicts that mechanical forces exerted on the cells synthesizing auxin; forces which would be sufficient to direct the development of vascular strands. Indeed, the importance of geometrical and mechanical constraints during vascular tissue development in the Arabidopsis embryonic root has already been underlined (De Rybel et al., 2014). Our data show that unlike *PIN1*, whose expression is clearly stimulated by auxin, biosynthetic genes are either insensitive to, or repressed by auxin. Moreover, i) TAA1 expression is reduced, not increased, as PIN1 accumulates and polarizes in provascular cells; ii) PIN1 expression is strongly down-regulated in leaves of auxin biosynthesis mutants; and iii) whilst TAA1 can be seen in domains in which PIN1 is absent, the opposite case is never observed. Taken together, these observations suggest that local auxin biosynthesis in leaf primordia is not part of the auxin canalization positive feedback loop and plays an integral role in leaf vasculature development.

So far our model only focuses on the development of the midvein. The initial branching stages to secondary and tertiary veins is likely to require more complex models, which are outside the scope of this work. The search for factors which drive vascular patterning needs urgently to be redirected to include the highly complicated patterns of auxin biosynthesis. The regulation of auxin biosynthesis by specific transcription factors (Cui et al., 2013) may complete such a mechanism.

## Author Contributions

F.A.D., W.D.T., I.K. and K.P. conceived and designed the experiments. F.A.D., I.K. performed the experiments. J.D. performed the mathematical modeling. F.A.D., I.K., W.D.T., E.K., R.T. and K.P. analyzed the data. F.A.D., W.D.T, E.K. and K.P. wrote the paper. All authors discussed the results and commented on the manuscript. Correspondence and request for material should be addressed to F.A.D. and K.P.

## Acknowledgements

This work could not have been accomplished without the help of colleagues, collaborators and friends who provided support, suggestions and materials. Particularly we would like to thank Jose Alonso and Yunde Zhao for sharing materials. We also gratefully acknowledge the excellent technical support from Beata Ditengou and Katja Rapp.

This work was supported by the Baden-Württemberg Stiftung, Deutsche Forschungsgemeinschaft (SFB 746), the Excellence Initiative of the German Federal and State Governments (EXC 294), Bundesministerium für Forschung und Technik (BMBF SYSTEC, PROBIOPA, MICROSYSTEMS), Deutsches Zentrum für Luft und Raumfahrt (DLR 50WB1022), the Freiburg Initiative for Systems Biology, the European Union Framework 6 Program (AUTOSCREEN, LSHG-CT-2007-037897), the National Science Foundation (USA), and JSPS KAKENHI (Grant Number 24570047, 16K07396). EK acknowledges support from the Burroughs Wellcome Fund.

## Methods

### Plant material and growth conditions

Columbia Arabidopsis ecotype (Col-0) as wild type (WT), *YUCp::GUS* reporter lines and *yuc* mutants single and multiple combinations were as previously described (Cheng et al., 2006; Ditengou et al., 2008). Details of *pTAA 1::TAA1-GFP, wei8-1, wei8-1tar2-1* and *wei8-1tar1-1tar2-1* and *pin1* mutants are as previously described (Stepanova et al., 2008); (Galweiler et al., 1998). *Wei8* (Stepanova et al. Cell 2008) is also known as *sav3* (Tao et al., 2008). To more accurately reflect the corresponding protein′s function, *WEI8* was subsequently renamed *TAA1 TRYPTOPHAN AMINOTRANSFERASE OF ARABIDOPSIS1* by Tao et al. (2008). YUC4p::GFP seeds were obtained from Yunde Zhao. Seeds were surface-sterilized and sown on solid Arabidopsis medium (2.3 g/liter MS salts, 1% sucrose, 1.6% agar-agar (pH 6.0) adjusted with KOH). After vernalization for 2 days at 4°C, seeds were germinated under a long day period (16h light, 8h darkness). In polar auxin transport inhibitory experiments, seedlings were grown on media supplemented with 10 μM NPA for 0 to 10 days.

### Immunocytodetection

Seedlings of different ages were fixed with 4% paraformaldehyde in PBS (pH 7.3) and used for whole-mount *in situ* immunolocalisation as previously described (Ditengou et al., 2008). PIN1 was detected using a mouse anti-PIN1 monoclonal antibody (1:100) and TAA1-GFP with a rabbit anti-GFP antibody (1:600) (Molecular Probes). YUC4p::GFP was detected with a rabbit anti-GFP polyclonal antibody (1:200). Samples were incubated with secondary antibodies (Alexa 488 goat anti mouse and Alexa 555 goat anti rabbit from Invitrogen, both at 1:1000 dilution).

### Microscopy and analysis

Histological detection of ·-glucuronidase (GUS) activity was performed according to (Scarpella et al., 2004). For analysis of vascular patterns, seedlings were cleared in 100% ethanol overnight. They were re-hydrated and dissected under 50% glycerol, then mounted in chloral hydrate:glycerol:water (8:3:1, w/v/v). GFP plants were fixed with 4% formaldehyde at room temperature and mounted in Prolong Gold antifade reagent (Molecular Probes). For light microscopy, samples were observed with a Zeiss Axiovert 200M MOT (Carl Zeiss, Goettingen, Germany) for high magnification pictures. By contrast, low magnification views were taken with a Zeiss Stemi SV11 Apo stereomicroscope (Carl Zeiss, Goettingen, Germany), viewed under differential interference contrast (DIC) optics or dark field illumination. Fluorescent proteins were analyzed with a Zeiss LSM 5 *DUO* scanning microscope or AZ-C1 Macro Laser Confocal Microscope (NIKON GMBH, Düsseldorf, Germany). GFP was excited using the 488nm laser line in conjunction with a 505-530 band-pass filter. To simultaneously monitor Alexa488 and Alexa 555 fluorescences, we used multi-tracking in frame mode and the emission was separated using the on-line unmixing feature of the Meta spectral analyzer. Images were extracted and analyzed with the Zen2009 software (Carl Zeiss MicroImaging) and representative of at least 20 individual plants.

### Real time RT-PCR

The effect of auxin on gene expression was quantified by real-time quantitative RT-PCR. Arabidopsis seeds were deposited on 6-7 mm filter paper strips lying at the surface of solid Arabidopsis growth medium (see above). This procedure facilitates the transfer of seedlings on the filter paper from Agar to liquid Arabidopsis growth medium (not containing Agar) supplemented with or without IAA. Three-day old Arabidopsis seedlings were transferred to medium containing either mock (control) or IAA. After treatment the filter paper strips with seedlings were transferred to RNA-later solution (Ambion). Total RNA was extracted using the RNeasy Micro Kit (Qiagen). Reverse transcription was performed using 1μg of total RNA and RevertAid M-MulV reverse transcriptase (Fermentas, St Leon-Rot, Germany) according to the manufacturer′s instructions. qRT-PCR was performed using Maxima SYBR Green kit (Fermentas) on a Light cycler480 Real-Time system (Roche, Mannheim, Germany). The gene-specific primers for qRT-PCR are listed in Table S2. The efficiency of each primer pair was determined by examination of a standard curve using serial dilutions of genomic DNA. The PCR was performed using a three-step protocol including melting curve analysis. The relative gene expression was analyzed using the Δ-Δ cycle threshold method. ACTIN2 served reference gene, three biological replicates and three technical replicates were used to evaluate gene expression.

### Image post-processing

All images depicting PIN1/TAA1 double labeling or PIN1/DAPI are 3D reconstructions of optical sections with Imaris 7.4.0 (Bitplane AG, Zurich, Switzerland). All images were assembled using Microsoft PowerPoint 2013.

### Computational model description

A cell-based modeling framework, Virtual Leaf, that couples vertex dynamics and chemical dynamics, was adapted to study the vein patterning in leaves *in siiico* (Merks et al., 2011). We simplified the model system by representing the longitudinal section of leaf primordia by a two-dimensional network of interconnected polygons (cells), specified by the surrounding vertices. Each cell, in this framework, is characterized by an energy function which describes the balance between turgor pressure and cell wall stiffness,

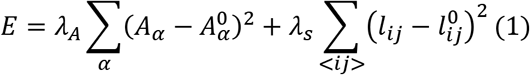

*A_α_* is the area of the cell *α* and 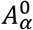 is the preferred (rest) area of that cell. *l_ij_* is the length of the cell wall linking nodes *i* and *j* of the polygons and the sum over < *ij* > is over all links. *λ_A_* is a parameter setting the cell resistance to compression or expansion and *λ_s_* describes the cell wall stiffness. Cell behaviors, like expansion, division and active shape changes are described by minimization of the energy function *E*. The minimization depends on the dimensionless ratio 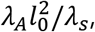, where *l_0_* is a cell size dependent lengthscale. The value of the dimensionless ratio used in our simulations was 0.01, in accordance with (Merks et al., 2011).

To study vein patterning in developing leaf primordia, we created a leaf template to resemble the tissue of a leaf primordium, as in Fig. 5A. Cells at the base of this template (cells colored in gray in Fig. 5A) were assigned with very high cell stiffness *λ_s_*, in order to implement proper boundary conditions, namely the effect of the firm attachment of the leaf primordia to the shoot apical meristem. The outermost layer of cells in the leaf primordia represents the stiff epidermal layer. To incorporate these properties of the epidermal cells in our model, the outermost cell walls of the cells at the perimeter of the leaf primordium were assumed to have higher stiffness than the internal cell walls. We used a value for the ratio of perimeter to internal cell wall stiffness within the experimentally reported range, i.e *λ_s_*(perimeter) = 12*λ_s_*(interior) (Gibson et al., 1988; Onoda et al., 2015). In our model auxin is synthesized locally in only few cells and subsequently auxin diffuses across the neighboring cells in the tissue. The leaf primordia in our model consists of three different cell types:

(1) auxin synthesizing cells (colored in dark green in Fig. 4A),
(2) cells that do not synthesize auxin (colored in light green in Fig. 4A),
(3) petiole cells (colored in grey in Fig. 4A).

The latter is a computational cell type aimed to simulate the firm attachment of the leaf to the meristem and the drainage of auxin.

The general dynamic equation governing the amount of auxin *n_α_* in a cell is given by:

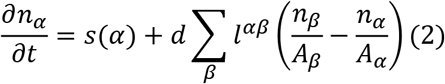

Here *l^αβ^* is the length of the cell wall separating cells *α* and *β*, across which the auxin diffusion takes place and *A_α_* is the area of the cell *α*. The first term describes auxin production at a constant rate *s(α) = s_0_δ_α,p_*, non-zero only for the auxin producer cells *p*. Auxin production is limited to only few cells (cell colored in dark green in Fig. 4A). The second term accounts for auxin spreading in the tissue of leaf primordia by a diffusion process, where *d* is a constant that measures the speed of inter-cellular auxin transport and is related to the auxin diffusion coefficient. Auxin is drained through cells located at the base of leaf primordia. To simulate this auxin drainage, from these cells (termed “petiole cells” and colored in grey in Fig. 4A), we implement perfectly absorbing boundary condition at the petiole cell walls.

The typical cell size in the simulation has a length of about *l_0_ ≅ 10μm*. We used the experimentally reported range, 10^−5^ - 10^−8^*ms*^-1^, for the value of speed of auxin transport *d* (Mitchison, 1980). At time *t* = 0 the amount of auxin is *n* = 0 in all the cells in the leaf primordia. For *t* > 0 auxin is produced only in auxin synthesizing cells at a constant rate *s*_0_. Motivated by experimental images of a 2DAG leaf, in our simulation we start initially with only four auxin synthesizing cells, but our qualitative results are not dependent on the initial geometry (Fig. 4A left and right panel and Fig. S7). All the cells in the leaf primordia grow by increasing their target area at the same constant rate *g*_0_ = 610^−14^*m*^2^*s*^-1^ and divide over their shortest axis once their area is doubled. However, in any cell except the auxin synthesizing cell, if the auxin concentration increases beyond a threshold value *c*\* = 2.410^9^*m*^-2^, then the growth rate of that cell increases to a value *g > g*_0_. We tested a broad range of cell area growth rates *g* and found that main vein formation and main vein bifurcation is a robust feature appearing for the values of ratio *g/g*_0_ ≳ 3 and up to at least several hundreds (Fig. S8).

The force *F(X_i_*) on each node *i* at vector position *X_i_* of a cell is given by:

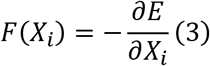

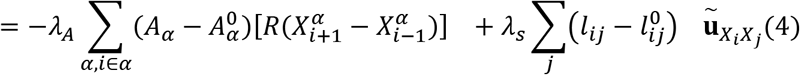

where *R* is the 90 degree rotation matrix:

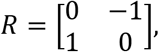

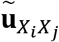 is the unit vector joining the vertices at position *X_i_* and *X_j_*, the summation *Σ_α,i∈a_* is over all cells that contain node *i* and 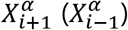 denotes the subsequent (antedecent) to node *i* as the nodes in cell *α* are traversed clockwise.

The force vector field of an *in silico* growing leaf primordium is shown by arrows in Fig. 4B,D,F. Magenta colored arrows represent net force acting on the vertices of auxin producing cells, whereas blue colored arrows represent net force acting on the vertices of non-auxin producing cells in the leaf primordia. The values for the model parameters are shown in Table S1.

### PIN1 based auxin transport

In addition to the passive diffusive transport of auxin, we also studied, in our model, the effect of active transport of auxin via PIN1 only. The general dynamic equation governing the amount of auxin *n_α_* in cell *α* in this case is given by:

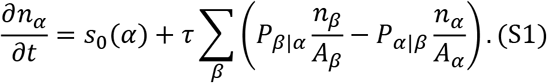

Here *P_β|α_* is the amount of PIN1 on the cell wall of cell *β* neighboring cell *α, τ* is the constant related to the auxin transport via PIN1 and *A_β_* is the area of cell *β*. The above equation is an extension of Equation (1) given above with *d* = 0. The only addition being the second term on the right hand side of Equation S1, which describes the intercellular transport of auxin via PINl.

The rate of attachment of PIN1 to a given cell wall that separates cell *α* and cell *β* depends on the amount of auxin in that particular cell, *α* and its neighboring cell*β*, as well as the amount of cytoplasmic PIN1, *P_α_* in that cell. The flux of PIN1 molecules attaching to a given cell wall is given by:

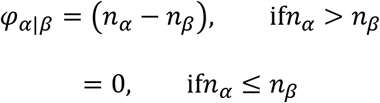

Equation governing the amount of PIN1 on the wall of a given cell is given by:

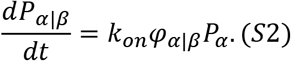

Here, *P_α_* is the amount of cytoplasmic PIN1 in cell *α* and *k_on_* is the rate of association of cytoplasmic PIN1 to the wall of cell *α* neighboring cell *β*.

The dynamic equation governing the amount of cytoplasmic PIN1, *P_α_* in cell *α* is given by:

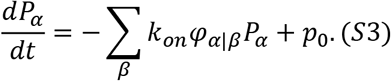

where *p*_0_ is the rate of PIN1 production in each and every cell.

Figure 6G shows the simulation result of our model including the PIN1 based auxin transport, described by Equations S1-S3, with the parameter values given in Table S3.

Our simulation results show the formation of a normal mid-vein also in the case of no diffusion (*d* = 0) and PIN1 only transport, see Figure 4G (main text).

### Compensation of vein patterning in auxin biosynthesis mutants by NPA

In our model, the area growth rate of non-auxin producing cells depend on the cellular auxin concentration. When the auxin concentration in a non-auxin producing cell crosses a specified threshold, its area growth rate increases, as a result these cells grow faster than the auxin producing cells, inducing transverse forces, acting laterally on the walls of midvein cells, much larger in magnitude than the forces acting on other cells in the leaf primordia. This prevents the midvein cells from proliferating, thus resulting in a thin vascular strand of elongated cells.

In this section, using our model, we will see how reducing auxin production rate requires a simultaneous reduction in auxin transport rate as well as overall cell area growth rate in order to recover wild-type midvein patterning. Our model therefore proposes a mechanism through which a reduction in auxin biosynthesis can rescue the NPA-induced defects caused by auxin accumulation in the midvein.

Continuous auxin production in auxin synthesizing cells and inter-cellular auxin transport sets up an auxin concentration gradient in the growing leaf primordia. Therefore, auxin concentration threshold, beyond which non-auxin producing cells start to grow faster, depends upon several kinetic parameters that also determine auxin concentration gradient.

Auxin concentration of a cell in a growing leaf lamina changes according to the following equation:

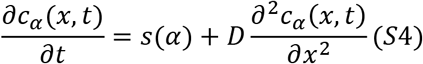

where, *c_α_(x,t*) is the auxin concentration of cell *α* at position *x* and time *t* in the leaf primordia, *s(α) - s_0_δ_α,p_*, describes auxin production at a constant rate *s*_0_, non-zero only for auxin producing cells *p*, and *D* is the auxin diffusion. Since leaf primordia is growing in area due to an irreversible growth in the area of an individual cell, as a result, the cellular auxin concentration *c = n/A*, where *n* is the number of auxin molecules in a cell of area *A*, is effected,

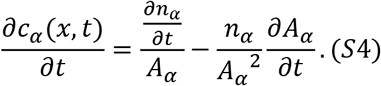

The first term in Equation (S4) describes the change in auxin concentration due to auxin transport, and is given by Equation (2) in the main text. The second term describes the change in auxin concentration as result of dilution of auxin due to an increase in cell area due to cell area growth. Similar situation of an area growth induced dilution of a chemical specie, growth hormone, morphogens, etc., is encountered in other biological systems (Romanova-Michaelides et al., 2015; Wartlick et al., 2011).

The change in auxin concentration of cell *α* at position *x* and time *t*, can thus be written down as:

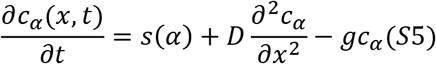

where, *g* is the area growth of a cell. The steady state solution of Equation (S5) for a delta source of auxin is given by:

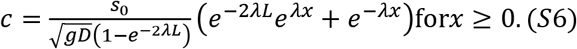

Equation (S6) shows the dependence of auxin concentration threshold *c*\*on auxin production rate *s*_0_, auxin diffusion rate *D*, and cell area growth rate *g*. Auxin diffusion rate is related to the auxin transport rate *d*, which is the actual parameter used in our model to describe auxin transport. It can be seen that lowering auxin productions_0_, as in auxin biosynthesis mutants, requires a simultaneous lowering of auxin transport rate *d*, similar to NPA treatment, as well as overall cell area growth rate *g* (Figures S7 and 4F, main text).

## REFERENCES

Abley, K., Sauret-Gueto, S., Maree, A.F.M., and Coen, E. (2016). Formation of polarity convergences underlying shoot outgrowths. Elife 5. https://doi.org/10.7554/eLife.18165.001.

Avsian-Kretchmer, O., Cheng, J.C., Chen, L.J., Moctezuma, E., and Sung, Z.R. (2002). Indole acetic acid distribution coincides with vascular differentiation pattern during Arabidopsis leaf ontogeny. Plant Physiology 130, 199–209. DOI: http://dx.doi.org/10.1104/pp.003228.

Bayer, E.M., Smith, R.S., Mandel, T., Nakayama, N., Sauer, M., Prusinkiewicz, P., and Kuhlemeier, C. (2009). Integration of transport-based models for phyllotaxis and midvein formation. Genes & Development 23, 373–384. DOI: 10.1101/gad.497009.

Blonder, B., Violle, C., Bentley, L.P., and Enquist, B.J. (2011). Venation networks and the origin of the leaf economics spectrum. Ecol Lett 14, 91–100. DOI: 10.1111/j.1461-0248.2010.01554.x.

Bruck, D.K., and Paolillo, D.J. (1984). Replacement of leaf primordia with IAA in the induction of vascular differentiation in the stem of Coculus. New Phytologist 96, 353–370.

Cai, X.T., Xu, P., Zhao, P.X., Liu, R., Yu, L.H., and Xiang, C.B. (2014). Arabidopsis ERF109 mediates crosstalk between jasmonic acid and auxin biosynthesis during lateral root formation. Nature Communications 5, 5833. DOI: 10.1038/ncomms6833.

Carland, F., Defries, A., Cutler, S., and Nelson, T. (2016). Novel Vein Patterns in Arabidopsis Induced by Small Molecules. Plant Physiology 170, 338–353. DOI: 10.1104/pp.15.01540.

Cheng, Y., Dai, X., and Zhao, Y. (2006). Auxin biosynthesis by the YUCCA flavin monooxygenases controls the formation of floral organs and vascular tissues in Arabidopsis. Genes Dev 20, 1790–1799. DOI: 10.1101/gad.1415106.

Cui D., Zhao J., Jing Y., Fan M., Liu J., Wang Z., Xin W., and Hu, Y. (2013). The arabidopsis IDD14, IDD15, and IDD16 cooperatively regulate lateral organ morphogenesis and gravitropism by promoting auxin biosynthesis and transport. PLoS Genet 9, e1003759. DOI: 10.1371/journal.pgen.1003759.

De Rybel, B., Adibi, M., Breda, A.S., Wendrich, J.R., Smit, M.E., Novak, O., Yamaguchi, N., Yoshida, S., Van Isterdael, G., Palovaara, J., et al. (2014). PLANT DEVELOPMENT Integration of growth and patterning during vascular tissue formation in Arabidopsis. Science 345. DOI:10.1126/science.1255215.

Ditengou, F.A., Teale, W.D., Kochersperger, P., Flittner, K.A., Kneuper, I., van der Graaff, E., Nziengui, H., Pinosa, F., Li, X., Nitschke, R., et al. (2008). Mechanical induction of lateral root initiation in Arabidopsis thaliana. Proceedings of the National Academy of Sciences of the United States of America 105, 18818–18823. DOI: 10.1073/pnas.0807814105.

Effendi, Y., Ferro, N., Labusch, C., Geisler, M., and Scherer, G.F. (2015). Complementation of the embryo-lethal T-DNA insertion mutant of AUXIN-BINDING-PROTEIN 1 (ABP1) with abp1 point mutated versions reveals crosstalk of ABP1 and phytochromes. Journal of Experimental Botany 66, 403–418. DOI: 10.1093/jxb/eru433.

Feller, C., Farcot, E., and Mazza, C. (2015). Self-Organization of Plant Vascular Systems: Claims and Counter-Claims about the Flux-Based Auxin Transport Model. PLoS One 10. DOI: 10.1371/journal.pone.0118238.

Fendrych, M., Leung, J., and Friml, J. (2016). TIR1/AFB-Aux/IAA auxin perception mediates rapid cell wall acidification and growth of Arabidopsis hypocotyls. Elife 5. DOI: 10.7554/eLife.19048.

Galweiler, L., Guan, C., Muller, A., Wisman, E., Mendgen, K., Yephremov, A., and Palme, K. (1998). Regulation of polar auxin transport by AtPIN1 in Arabidopsis vascular tissue. Science 282, 2226–2230.

Gibson, L.J., Ashby, M.F., and Easterling, K.E. (1988). Structure and Mechanics of the Iris Leaf. J Mater Sci 23, 3041–3048. DOI: 10.1016/j.jbiomech.2004.09.027.

Hamant, O., Heisler, M.G., Jonsson, H., Krupinski, P., Uyttewaal, M., Bokov, P., Corson, F., Sahlin, P., Boudaoud, A., Meyerowitz, E.M., et al. (2008). Developmental Patterning by Mechanical Signals in Arabidopsis. Science (Washington D C) 322, 1650–1655. DOI: 10.1126/science.1165594.

Heisler, M.G., Ohno, C., Das, P., Sieber, P., Reddy, G.V., Long, J.A., and Meyerowitz, E.M. (2005). Patterns of auxin transport and gene expression during primordium development revealed by live imaging of the Arabidopsis inflorescence meristem. Curr Biol 15, 1899–1911. DOI: 10.1016/j.cub.2005.09.052

Himanen, K., Boucheron, E., Vanneste, S., Engler, J.D., Inze, D., and Beeckman, T. (2002). Auxin-mediated cell cycle activation during early lateral root initiation. Plant Cell 14, 2339–2351. DOI: http://dx.doi.org/10.1105/tpc.004960.

Ichihashi, Y., Kawade, K., Usami, T., Horiguchi, G., Takahashi, T., and Tsukaya, H. (2011). Key Proliferative Activity in the Junction between the Leaf Blade and Leaf Petiole of Arabidopsis. Plant Physiology 157, 1151–1162. DOI: http://dx.doi.org/10.1104/pp.111.185066.

Keller, C.P., Stahlberg, R., Barkawi, L.S., and Cohen, J.D. (2004). Long-term inhibition by auxin of leaf blade expansion in bean and arabidopsis. Plant Physiology 134, 1217–1226. doi: 10.1104/pp.103.032300.

Lee, S.-W., Feugier, F.G., and Morishita, Y. (2014). Canalization-based vein formation in a growing leaf. J Theor Biol 353, 104–120. DOI: 10.1016/j.jtbi.2014.03.005.

Mähönen, A.P., ten Tusscher, K., Siligato, R., Smetana, O., Díaz-Triviño, S., Salojärvi, J., Wachsman, G., Prasad, K., Heidstra, R., and Scheres, B. (2014). PLETHORA gradient formation mechanism separates auxin responses. Nature 515, 125–129. DOI: 10.1038/nature13663.

Mashiguchi, K., Tanaka, K., Sakai, T., Sugawara, S., Kawaide, H., Natsume, M., Hanada, A., Yaeno, T., Shirasu, K., Yao, H., et al. (2011). The main auxin biosynthesis pathway in Arabidopsis. Proceedings of the National Academy of Sciences of the United States of America 108, 18512–18517. DOI: 10.1073/pnas.1108434108.

Mattsson, J., Ckurshumova, W., and Berleth, T. (2003). Auxin signaling in Arabidopsis leaf vascular development. Plant Physiology 131, 1327–1339.

Mattsson, J., Sung, Z.R., and Berleth, T. (1999). Responses of plant vascular systems to auxin transport inhibition. Development 126, 2979–2991.

Merks, R.M.H., Guravage, M., Inzé, D., and Beemster, G.T.S. (2011). VirtualLeaf: an open-source framework for cell-based modeling of plant tissue growth and development. Plant Physiol 155, 656–666. DOI: 10.1104/pp.110.167619.

Mitchison G.J. (1980). The Dynamics of Auxin Transport, Vol 209.

Mitchison G.J. (1981). The Polar Transport of Auxin and Vein Patterns in Plants. Philos T Roy Soc B 295, 461–471. DOI: 10.1098/rstb.1981.0154.

Nishimura, T., Hayashi, K., Suzuki, H., Gyohda, A., Takaoka, C., Sakaguchi, Y., Matsumoto, S., Kasahara, H., Sakai, T., Kato, J., et al. (2014). Yucasin is a potent inhibitor of YUCCA, a key enzyme in auxin biosynthesis. Plant Journal 77, 352–366. DOI: 10.1111/tpj.12399.

Okada, K., Ueda, J., Komaki, M.K., Bell, C.J., and Shimura, Y. (1991). Requirement of the Auxin Polar Transport System in Early Stages of Arabidopsis Floral Bud Formation. The Plant Cell Online 3, 677–684. DOI: 10.1105/tpc.3.7.677.

Onoda, Y., Schieving, F., and Anten, N.P.R. (2015). A novel method of measuring leaf epidermis and mesophyll stiffness shows the ubiquitous nature of the sandwich structure of leaf laminas in broad-leaved angiosperm species. J Exp Bot 66, 2487–2499. DOI: 10.1093/jxb/erv024.

Pacheco-Villalobos, D., Diaz-Moreno, S.M., van der Schuren, A., Tamaki, T., Kang, Y.H., Gujas, B., Novak, O., Jaspert, N., Li, Z.N., Wolf, S., et al. (2016). The Effects of High Steady State Auxin Levels on Root Cell Elongation in Brachypodium. Plant Cell 28, 1009–1024. DOI: 10.1105/tpc.15.01057.

Perrot-Rechenmann, C. (2010). Cellular Responses to Auxin: Division versus Expansion. Cold Spring Harbor Perspectives in Biology 2. DOI: 10.1101/cshperspect.a001446.

Petersson, S.V., Johansson, A.I., Kowalczyk, M., Makoveychuk, A., Wang, J.Y., Moritz, T., Grebe, M., Benfey, P.N., Sandberg, G., and Ljung, K. (2009). An Auxin Gradient and Maximum in the Arabidopsis Root Apex Shown by High-Resolution Cell-Specific Analysis of IAA Distribution and Synthesis. Plant Cell 21, 1659–1668. DOI: 10.1105/tpc.109.066480.

Petrasek, J., Mravec, J., Bouchard, R., Blakeslee, J.J., Abas, M., Seifertova, D., Wisniewska, J., Tadele, Z., Kubes, M., Covanova, M., et al. (2006). PIN proteins perform a rate-limiting function in cellular auxin efflux. Science 312, 914–918. DOI: 10.1126/science.1123542.

Rolland-Lagan, A.G., and Prusinkiewicz, P. (2005). Reviewing models of auxin canalization in the context of leaf vein pattern formation in Arabidopsis. Plant Journal 44, 854–865. DOI:10.1111/j.1365-313X.2005.02581.x.

Romanova-Michaelides, M., Aguilar-Hidalgo, D., Julicher, F., and Gonzalez-Gaitan, M. (2015). The wing and the eye: a parsimonious theory for scaling and growth control? Wiley Interdiscip Rev Dev Biol. 4, 591–608. DOI: 10.1002/wdev.195.

Tao, Y., Ferrer, J.L., Ljung, K., Pojer, F., Hong, F., Long, J.A., Li, L., Moreno, J.E., Bowman, M.E., Ivans, L.J., et al. (2008). Rapid synthesis of auxin via a new tryptophan-dependent pathway is required for shade avoidance in plants. Cell 133, 164–176. doi: 10.1016/j.cell.2008.01.049.

Sachs T. (1969). Polarity and Induction of Organized Vascular Tissues. Annals of Botany 33, 263–275.

Sampathkumar, A., Yan, A., Krupinski, P., and Meyerowitz, E.M. (2014). Physical Forces Regulate Plant Development and Morphogenesis. Current Biology 24, 475–R483. DOI: 10.1016/j.cub.2014.03.014.

Sawchuk, M.G., Edgar, A., and Scarpella, E. (2013). Patterning of Leaf Vein Networks by Convergent Auxin Transport Pathways. PLoS Genetics 9. DOI: 10.1371/journal.pgen.1003294.

Scarpella, E., Francis, P., and Berleth, T. (2004). Stage-specific markers define early steps of procambium development in Arabidopsis leaves and correlate termination of vein formation with mesophyll differentiation. Development 131, 3445–3455. DOI:10.1242/dev.01182.

Scarpella, E., Marcos, D., Friml, J., and Berleth, T. (2006). Control of leaf vascular patterning by polar auxin transport. Genes Dev 20, 1015–1027. DOI:10.1101/gad.1402406.

Scheres B., and Xu, L. (2006). Polar auxin transport and patterning: grow with the flow. Genes & Development 20, 922–926. DOI:10.1101/gad.1426606.

Sieburth L.E. (1999). Auxin is required for leaf vein pattern in Arabidopsis. Plant Physiology 121, 1179–1190. DOI: http://dx.doi.org/10.1104/pp.121.4.1179.

Stepanova, A.N., Robertson-Hoyt, J., Yun, J., Benavente, L.M., Xie, D.Y., Dolezal, K., Schlereth, A., Jurgens, G., and Alonso, J.M. (2008). TAA1-mediated auxin biosynthesis is essential for hormone crosstalk and plant development. Cell 133, 177–191. DOI: 10.1016/j.cell.2008.01.047.

Stepanova, A.N., Yun, J., Robles, L.M., Novak, O., He, W., Guo, H., Ljung, K., and Alonso, J.M. (2011). The Arabidopsis YUCCA1 Flavin Monooxygenase Functions in the Indole-3-Pyruvic Acid Branch of Auxin Biosynthesis. Plant Cell 23, 3961–3973. DOI: 10.1105/tpc.111.088047.

Thomson, K.S., Hertel, R., Muller, S., and Tavares, J.E. (1973). 1-N-Naphthylphthalamic Acid and 2,3,5-Triiodobenzoic Acid - in-Vitro Binding to Particulate Cell Fractions and Action on Auxin Transport in Corn Coleoptiles. Planta 109, 337–352. DOI: 10.1007/BF00387102.

Tsugeki, R., Ditengou, F.A., Sumi, Y., Teale, W., Palme, K., and Okada, K. (2009). NO VEIN Mediates Auxin-Dependent Specification and Patterning in the Arabidopsis Embryo, Shoot, and Root. Plant Cell 21, 3133–3151. DOI: 10.1105/tpc.109.068841.

Wartlick, O., Mumcu, P., Julicher, F., and Gonzalez-Gaitan, M. (2011). Understanding morphogenetic growth control - lessons from flies. Nat Rev Mol Cell Bio 12, 594–604. doi: 10.1038/nrm3169.

Won, C., Shen, X., Mashiguchi, K., Zheng, Z., Dai, X., Cheng, Y., Kasahara, H., Kamiya, Y., Chory, J., and Zhao, Y. (2011). Conversion of tryptophan to indole-3-acetic acid by TRYPTOPHAN AMINOTRANSFERASES OF ARABIDOPSIS and YUCCAs in Arabidopsis. Proceedings of the National Academy of Sciences of the United States of America 108, 18518–18523. DOI: 10.1073/pnas.1108436108

Zhao Y. (2012). Auxin biosynthesis: a simple two-step pathway converts tryptophan to indole-3-acetic acid in plants. Mol Plant 5, 334–338. DOI: 10.1093/mp/ssr104.

